# Modbamtools: Analysis of single-molecule epigenetic data for long-range profiling, heterogeneity, and clustering

**DOI:** 10.1101/2022.07.07.499188

**Authors:** Roham Razaghi, Paul W. Hook, Shujun Ou, Michael C. Schatz, Kasper D. Hansen, Miten Jain, Winston Timp

## Abstract

The advent of long-read sequencing methods provides new opportunities for profiling the epigenome - especially as the methylation signature comes for “free” when native DNA is sequenced on either Oxford Nanopore or Pacific Biosciences instruments. However, we lack tools to visualize and analyze data generated from these new sources. Recent efforts from the GA4GH consortium have standardized methods to encode modification location and probabilities in the BAM format. Leveraging this standard format, we developed a technology-agnostic tool, modbamtools to visualize, manipulate and compare base modification/methylation data in a fast and robust way. modbamtools can produce high quality, interactive, and publication-ready visualizations as well as provide modules for downstream analysis of base modifications. Modbamtools comprehensive manual and tutorial can be found at https://rrazaghi.github.io/modbamtools/.

## Introduction

Direct single-molecule sequencing methods, e.g. Pacific Biosciences (PacBio) and Oxford Nanopore Technologies (ONT), have recently greatly expanded in throughput and yield. In addition to the canonical base sequencing data that these platforms generate, modifications on the nucleic acids can be measured directly, either via delays in the incorporation of bases (IPD, PacBio (Flusberg et al. 2010)) or perturbations in the electrical current (ONT (Simpson et al. 2017)). These have been accompanied by development of software tools to measure and call modifications within this data, but the output formats of these calls were not standardized precluding easy downstream development. Modification data files have typically been stored as enormous (terabyte scale) tsv/csvs and early efforts to incorporate 5-methylcytosine information from ONT into a “bisulfite-like” BAM file required complex manipulations(Lee et al. 2020).

More recently, the Global Alliance for Genomics and Health (GA4GH) (Rehm et al. 2021) standards group proposed an addition to the BAM file spec, incorporating two new tags (MM and ML) for SAM/BAM alignment files. The MM tag is used to locate the strand and position the modification was observed on, and the ML tag is the probability of each modification being present (http://samtools.github.io/hts-specs). Although these tags were introduced as an adaptation to long-read base modification data, it is anticipated that all technologies will eventually incorporate this file format.

Single molecule base modification callers have rapidly adapted to the new standard format. Currently, for nanopore data, most modification calling tools can output BAM files with tags, including guppy, bonito, Megalodon, and nanopolish (Simpson et al. 2017). Similarly, Primrose, and ccsmeth (Ni et al. 2022) can be used for PacBio reads. An updated list of compatible tools generating these alignment files can be found at https://rrazaghi.github.io/modbamtools/.

Here we introduce modbamtools, a suite of tools to explore modifications in single-molecule data using this new format. With this tool we generate interactive and batch visualization and analysis for methylation frequency and single-molecule methylation. Profiling methylation across individual molecules, we can look at coordination of long-range methylation effects, e.g. enhancer-promoter interactions, and the degree of variation of methylation “noise” within regions. We have also generated modules to phase reads by using genetic variation or through methylation alone via a read clustering approach, to enable exploration of allele-specific methylation and epigenetic heterogeneity.

## Results

### Usage and Examples

We developed modbamtools, a software package that provides analysis and interactive visualization of single-read base modification data along with other highly used formats for genomic tracks (GTF, bigwig, bedgraph, etc). Modbamtools utilizes core python modules including numpy (van der Walt et al. 2011), pandas (McKinney and Others 2011), scikit-learn (Pedregosa et al. 2011), pysam (Heger et al. 2014), click, plotly (Plotly Technologies Inc., 2015), modbampy, pybigwig (Ryan et al. 2016), pypdf2, pillow, and hdbscan (McInnes et al. 2017). We have made modbamtools easily accessible through PyPI (‘pip install modbamtools’).

The tool has three main elements (‘calcMeth’, ‘calcHet’, ‘cluster’) and a plotting function that allows for interactive plotting of single-read base modification data. This generates a multi-panel plot **(Figure 1)** consisting of an annotation track, methylation frequency track, and single-read plots. The annotation track can display other sets of genomics data including gene models, other epigenetic data (e.g. ENCODE ChIP-seq), and genetic variation. Methylation frequencies along with a smoothed average frequency is plotted on top of the reads similar to a conventional genome browser. The methylation frequency plot shows the per locus frequency of modified to total called bases. Finally, the single-read plots represent each individual single molecule with base modifications indicated as blue for unmodified and red for modified. These figures can be output as HTML, PDF, PNG, or SVG. The HTML provided is generated with plotly and is interactive, allowing magnification. Multiple plots can be output in batch mode by providing a BED file of regions of interest resulting in a multiple page HTML or PDF report.

**Figure 1:**
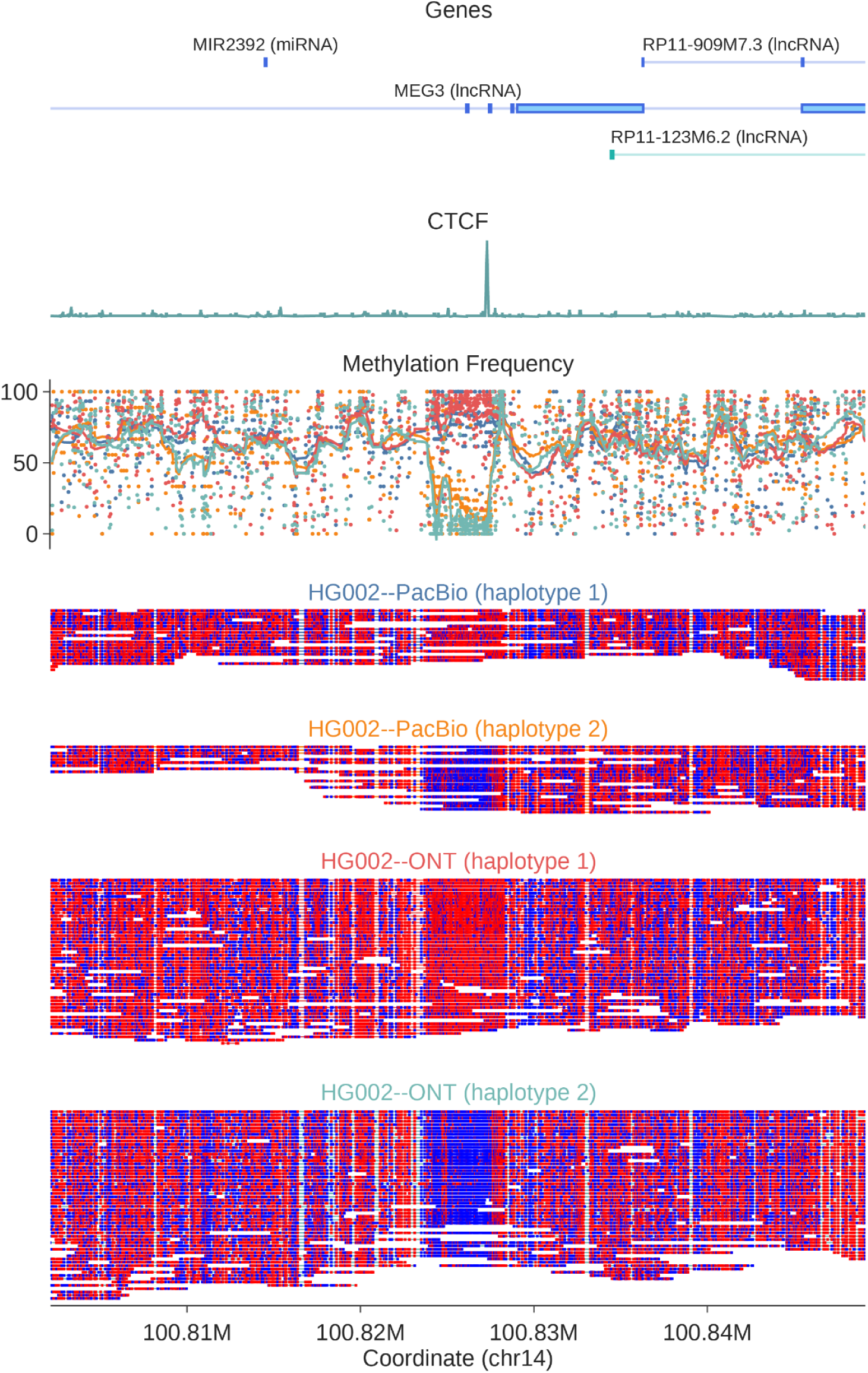
Example of modbamtools output on MEG3 (chr14:100,802,132-100,849,111) locus using both PacBio and ONT single-molecule data from the HG002 Genome in a Bottle cell line. “Genes” track shows GENCODE (Release 38, GRCH38) gene models and the “CTCF” track shows CTCF CHiP-seq ENCODE track from GM12878. Methylation frequency track is colored according to platform and haplotype, with colors indicated by the title of the single-molecule plots. In single-molecule plots, each read is a single horizontal bar, with methylated bases shown as red and unmethylated as blue.

Using appropriate tools, e.g. clair (Zheng et al. 2021) or whatshap (Martin et al. 2016), BAM files can have the haplotypes of reads encoded with the commonly used “HP” tag. Our tool has the ability to group the alignments based on phase tag (HP) in BAM files. Using this HP tag, we can separate reads according to haplotype, plotting each haplotype’s methylation frequency as different colored lines and the single reads as separate plot elements. We show an example of this module on methylation calls from the HG002 cell line at the *MEG3* long noncoding RNA (lncRNA), using public single-molecule methylation data from both ONT and PacBio platforms (**Figure 1**). *MEG3* has known monoallelic expression in many tissues and loss of this regulation has been implicated in development of type 2 diabetes mellitus (Rosa et al. 2005; Kameswaran et al. 2014). From this data, we observe clear examples of allele-specific methylation at a CTCF binding site and *MEG3* promoter region.

Beyond clustering according to genomic haplotype, we have implemented a method to cluster single-molecule reads based on methylation status alone using Hierarchical Density-Based Spatial Clustering of Applications with Noise (HDBSCAN) (McInnes et al. 2017). This is a useful feature for regions without many SNPs for phasing reads into haplotypes (Gershman et al. 2022). Clustering can also be used to quantify different cell types or to profile early cancer detection from a heterogeneous sample (Wang et al. 2021; Houseman et al. 2008; Gkountela et al. 2019; Tian et al. 2020). Clustering can be performed either as a part of the plotting command or separately (‘--cluster’ command) with the input of a batch file for locations used for the clustering. As shown in **Figure 2**, we can cluster the *SNURF* gene promoter based purely on methylation signal at this locus. This paternally imprinted locus can also be phased based on genotyping information (**Supplementary Figure 1**), demonstrating the agreement of our clustering approach with classical methods.

**Figure 2:**
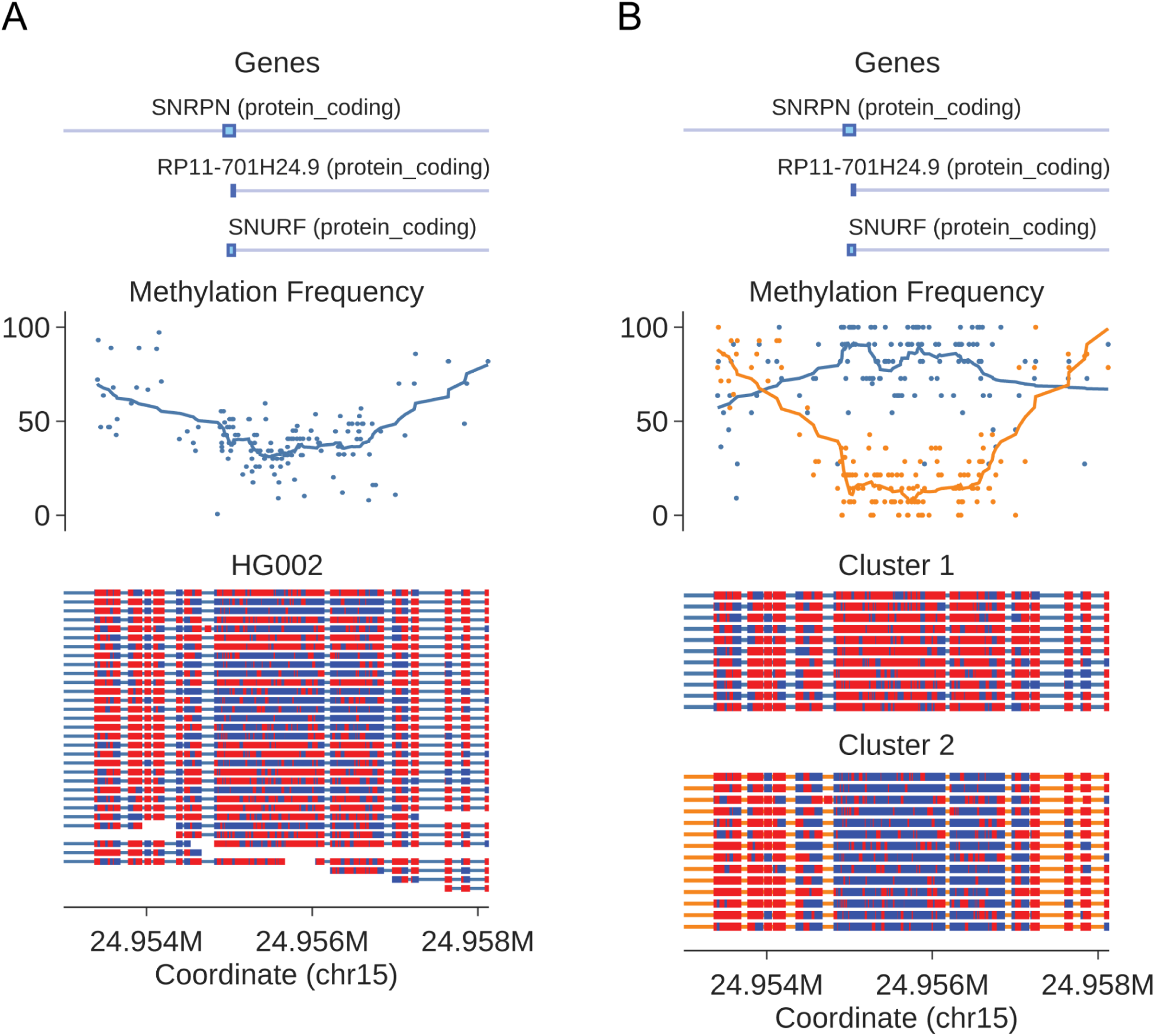
A) single molecule methylation profile on gene SNRPN (chr15:24,953,000-24,958,133) from HG002 data as in Figure 1. B) Single molecule methylation profile on gene SNRPN separated into clusters with ‘modbamtools plot –cluster’

Finally, using a BED file of genomic loci, we can profile the average methylation in each location, including methylation on each haplotype. The “calcMeth” module calculates methylation average across each single molecule *first* then aggregates over all molecules which map to that region, rather than averaging CpG methylation per CpG then averaging across the region. This is especially useful with long reads to capture methylation variability more efficiently (**Figure 3**).

**Figure 3:**
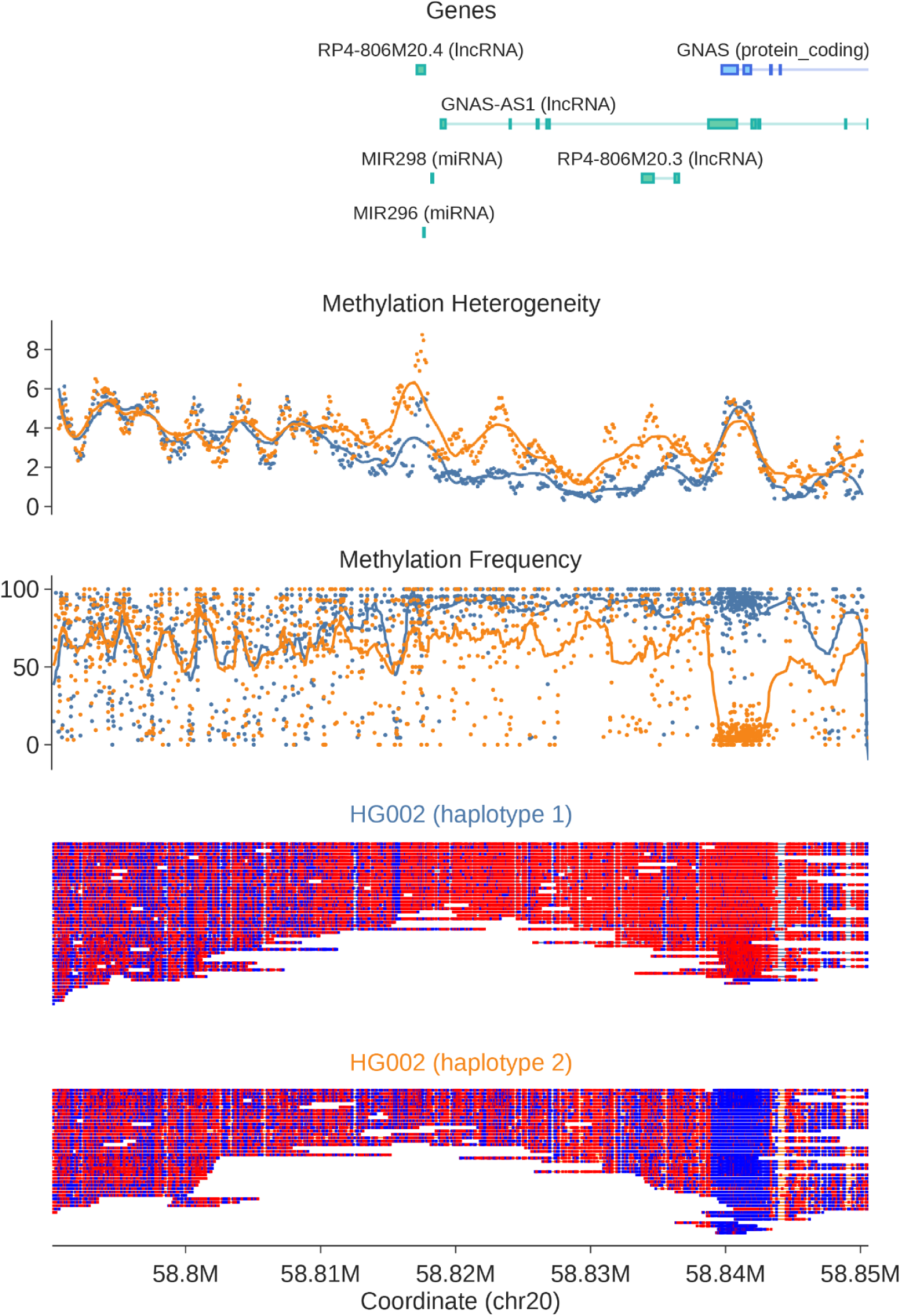
Example of modbamtools plot with options for haplotype separation and calculating heterogeneity at the GNAS locus (chr20:58,790,127-58,850,596).

With single-molecule methylation data, can quantify not only methylation frequency averaged across all reads, but also variability of methylation across individual molecules. A few studies have attempted to address this by proposing different algorithms to quantify this feature (Scherer et al. 2020; Landau et al. 2014; Guo et al. 2017; Landan et al. 2012; Xie et al. 2011). Here, we implemented a module to calculate methylation heterogeneity (“calcHet”) that calculates this on genomic regions provided by the user (See **Supplementary Note 1** for detailed methods). Similar to the clustering function, “–heterogeneity” option can be used with plotting command to visualize this; we have plotted it for the *GNAS* locus in **Figure 3**. There we observe areas of clear difference in methylation heterogeneity across the region, suggesting not only a change in methylation but a less ordered epigenetic state on one allele when compared to the other.

## Conclusion

Advances in single-molecule sequencing throughput suggest we are at an inflection point where large scale data sets are on the horizon. These data types offer the unique advantage of providing DNA methylation data *as well as* primary sequence - but without tools to take advantage of it, these data will be “left on the table” and not used to their potential. Here we have described a toolset to take advantage of these data, using the newly described modification tags present in the SAM/BAM file specifications. This toolset is compatible with all modern modification callers. Modbamtools provides fast, robust, interactive visualization and analysis for alignment files containing base modification tags.

## Acknowledgments

We would like to thank Jared Simpson and Chris Wright for their helpful comments and contributions to the development of modified base alignment files. W.T. has two patents (8,748,091 and 8,394,584) licensed to ONT.

## Data Availability

Publicly available data on cell line HG002 was downloaded from s3://ont-open-data/gm24385_mod_2021.09/extra_analysis/bonito_remora (ONT HG002 WGS) and https://downloads.pacbcloud.com/public/dataset/HG002-CpG-methylation-202202/ (PacBio HG002 WGS). CTCF track was downloaded from https://www.encodeproject.org/experiments/ENCSR000DZN/. GENCODE Release 38 for GRCH38 was used for gene model tracks.

## Code Availability and implementation

modbamtools source code is available at https://github.com/rrazaghi/modbamtools. A manual and tutorial are available at https://rrazaghi.github.io/modbamtools/.

