## Supplementary Information for "Modbamtools: Analysis of single-molecule epigenetic data for long-range profiling, heterogeneity, and clustering"

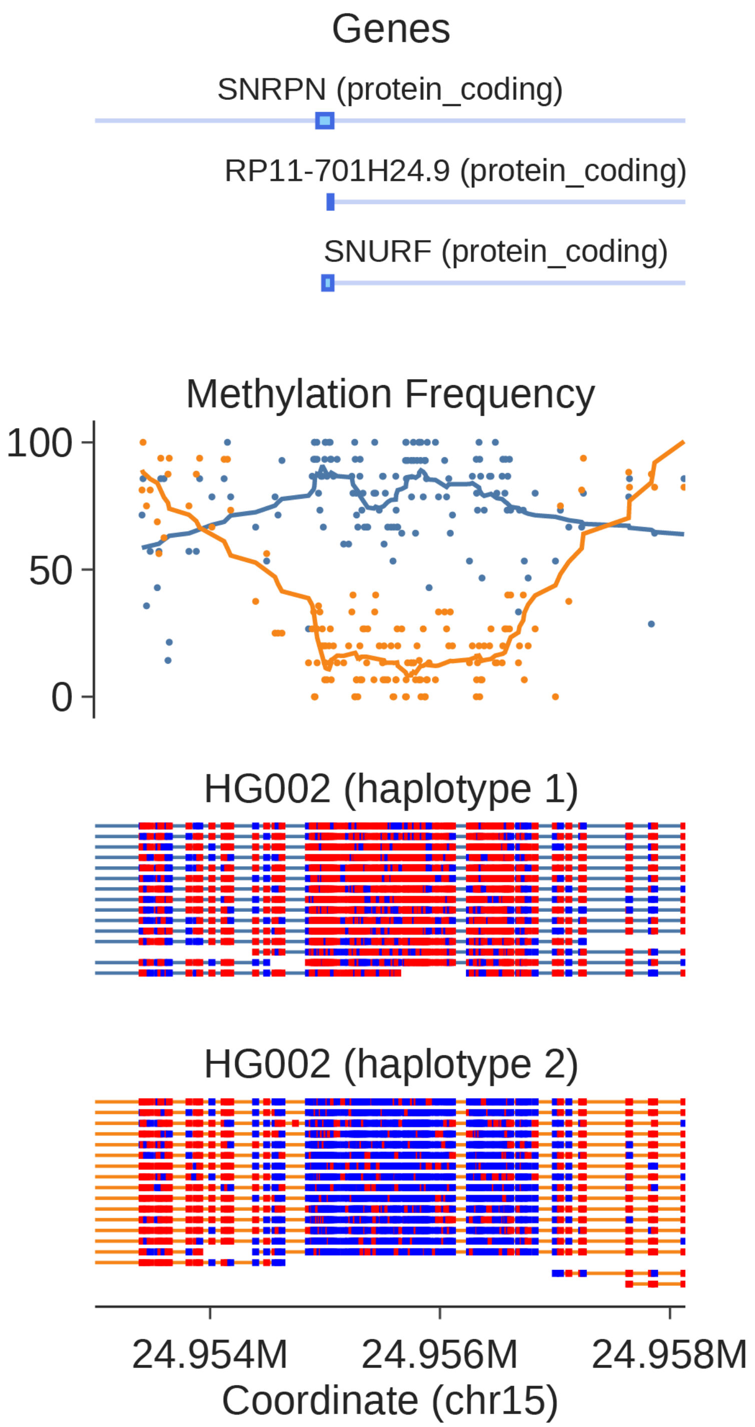


*Supplementary Figure 1: Single molecule methylation profile on gene SNRPN (chr15:24,953,000-24,958,133) from HG002 data as in Figure 2A separated by haplotypes*

**Supplementary Note 1:**

In order to calculate heterogeneity, we calculated the number of switches between runs of modified bases and unmodified bases. To compensate for possible sequencing/methylation calling error rates, we ignored singletons, i.e. runs consisting only an individual base. This was calculated using a 1kb bin side with a rolling window step size of 100bp. Average heterogeneity is calculated for each bin and smoothing is applied as in methylation frequency plots.
